# Accel-Align: A Fast Sequence Mapper and Aligner Based on the Seed–Embed–Extend Method

**DOI:** 10.1101/2020.07.20.211888

**Authors:** Yiqing Yan, Nimisha Chaturvedi, Raja Appuswamy

## Abstract

**Background:** Improvements in sequencing technology continue to drive sequencing cost towards $100 per genome. However, mapping sequenced data to a reference genome remains a computationally-intensive task due to the dependence on edit distance for dealing with indels and mismatches introduced by sequencing. All modern aligners use seed–filter–extend (SFE) methodology and rely on filtration heuristics to reduce the overhead of edit distance computation. However, filtering has inherent performance–accuracy trade-offs that limits its effectiveness.

**Results:** Motivated by algorithmic advances in randomized low-distortion embedding, we introduce *seed– embed–extend* (SEE), a new methodology for developing sequence mappers and aligners. While SFE focuses on eliminating sub-optimal candidates, SEE focuses instead on identifying optimal candidates. To do so, SEE transforms the read and reference strings from edit distance regime to the Hamming regime by embedding them using a randomized algorithm, and uses Hamming distance over the embedded set to identify optimal candidates. To show that SEE performs well in practice, we present Accel-Align, an SEE-based short-read sequence mapper and aligner that is 3-12*×* faster than state-of-the-art aligners on commodity CPUs, without any special-purpose hardware, while providing comparable accuracy.

**Conclusions:** As sequencing technologies continue to increase read length while improving throughput and accuracy, we believe that randomized embeddings open up new avenues for optimization that cannot be achieved by using edit distance. Thus, the techniques presented in this paper have a much broader scope as they can be used for other applications like graph alignment, multiple sequence alignment, and sequence assembly.

**Availability:** https://github.com/raja-appuswamy/accel-align-release

## 1 Introduction

Over the last decade, DNA sequencing technology has achieved dramatic improvements in both cost and throughput. With the $100-per-genome sequencing goal emerging as a realistic target in the near future, the amount of genomic data generated by sequencing is only poised to grow faster. The first, and often one of the most time consuming steps, in analyzing genomic datasets is sequence alignment–the process of determining the location in the reference genome of each sequencing read. Sequence alignment can be boiled down to a string matching problem. Given a string *G* as the reference genome, and a set of substrings *R* as the sequencing reads, the task of read alignment is to find the origin location of each read *r ϵ R* in *G*. However, due to sequencing errors, and differences between the reference genome and the sequenced organism, a read might not exactly align at any candidate location in the reference genome. Thus, an aligner has to perform approximate string matching to tolerate errors.

The error-handling capability of a string matching algorithm is related to the distance metric used for comparing strings. The two popular string distance metrics are Hamming distance which only counts the number of position-wise mismatches between two strings, and edit distance or Levenshtein distance, which counts the difference between two strings while allowing for characters to be inserted or deleted. Aligners can be classified into *ungapped* or *gapped* depending on whether they use Hamming or Levenshtein distance for computing mismatch between reads and the reference (Canzar and Salzberg (2017)). As modern sequencers can produce both substitution and indel errors, gapped aligners are preferred in practice over their ungapped counterparts.

Modern gapped read aligners, like Bowtie2, BWA-MEM, and Minimap2, can map thousands of reads per second to the reference genome. However, as sequencing datasets continue to grow at a rapid pace, even these state-of-the-art aligners face scalability bottlenecks due to a crucial design aspect that is universal across all gapped read aligners– the use of edit-distance as a string comparison metric. Computing edit distance between two sequences is a computationally-expensive task that takes approximately quadratic time in the length of the input sequences. Given that sub-quadratic computation of edit distance is extremely unlikely (Backurs and Indyk (2015)), the brute force approach of trying to align a read at each position in the reference is infeasible even for a single read due to sheer number of edit-distance computations that would be required. State-of-the-art aligners add to this complexity through the use of affine-gap penalty scoring function with optional soft clipping. Thus, all modern aligners focus on minimizing the number of such computations using a *seed-filter-extend* (SFE) strategy for performing alignment (Canzar and Salzberg (2017)).

SFE strategy works by first indexing the reference genome. Each read is processed in three steps. First, reads are broken down into smaller segments, referred to as *seeds*, and these seeds are used to look up potential mapping locations in the reference genome using an index. Second, candidate filtering techniques are used to filter out as many candidate locations as possible to minimize the overhead of the extension stage. Third, during the extension stage, the entire read is aligned at each of unfiltered candidate locations using the edit-distance-based approximate string matching algorithms.

Candidate filtering plays an important role in determining the overall throughput and scalability of read alignment, as it can eliminate many locations that would result in an incorrect mapping. However, current candidate filtering techniques present a performance–accuracy trade off. State-of-the-art candidate filtering techniques can be classified broadly as elimination-based or selection-based depending on the type of filtering strategy used. Elimination-based techniques, like adjacency filtering (Xin *et al*. (2013)), shifted hamming distance (Xin *et al*. (2015b)), focus on maintaining a high accuracy by conservatively eliminating candidate locations without significantly increasing the probability of misalignment due to accidental elimination of a true match. Recent research has demonstrated that such techniques create computational bottlenecks of their own that need to be addressed using specialized hardware (Kim *et al*. (2018); Alser *et al*. (2019)). Selection-based techniques, like voting in Subread (Liao *et al*. (2013b)), in contrast, use computationally less intensive selection criteria to directly pick a few candidate locations. Thus, they trade off accuracy for performance, as the selection criteria can end up eliminating a true match.

In this paper, we investigate a new selection-based candidate filtering strategy based on recent theoretical advances in the design of randomized algorithms that can perform embedding from edit distance into Hamming distance with very low distortion (Chakraborty *et al*. (2016)). These algorithms provide a one-to-one mapping *f* that can be used to transform a set of strings *S* into another set of strings *S’*, such that the worst case ratio between Hamming distance of any two strings *f(x)* and *f(y)* in *S’*, and the edit distance of their equivalent strings *x* and *y* in *S*, also called *distortion*, is very low. In this work, we investigate the use of such randomized algorithms in designing a *Seed-Embed-Extend* (SEE) aligner. An SEE aligner uses seeding to identify candidate locations similar to other SFE aligners. Randomized algorithms are then used to embed the reference string at each candidate location, and the Hamming distance between the embedded reference and the embedded read is used to rank the candidate locations based on likelihood of being an actual alignment target. Finally, candidates with the highest rank are chosen for extension.

To show that SEE works well in practice, we present *Accel-Align*– an SEE-based short-read sequence mapper and aligner that can provide both extension-free mapping and base-to-base alignment with CIGAR and MAPQ. In doing so, we show that a naive implementation of SEE will result in the embedding step becoming a computational bottleneck, and describe several optimizations that Accel-Align uses to implement SEE-based alignment effectively. Using experimental results from both simulated and real datasets, we show that embedding is capable of picking locations that are likely to be the correct alignment targets with very high accuracy. Using the SEE-approach to sequence alignment, Accel-Align can align 280,000 100bp reads per second on a commodity quad-core CPU, and is up to 9*×* faster than BWA-MEM, 12*×* faster than Bowtie2, and 3*×* faster than Minimap2, while providing comparable accuracy without using any special purpose hardware. We believe that SEE specifically, and embedding in general, is a robust technique that opens up new optimization opportunities not only for sequencing alignment, but also for several other computational biology problems that rely on edit distance.

## 2 Methods

Accel-Align adopts the SEE methodology for sequence alignment. During alignment, Accel-Align processes each read by first extracting seeds to find candidate locations. After seeding, Accel-Align embeds both the read string and the reference string at each candidate location found by seeding. After embedding, Accel-Align calculates the Hamming distance between each embedded reference and the read, and selects the two best candidate locations with the lowest Hamming distance for extension and scoring. In the rest of this section, we will introduce each of the three stages of SEE, and their implementation in Accel-Align, in detail.

### 2.1 Building the index and seeding

#### 2.1.1 Building the index

As Accel-Align uses seeding, it requires the reference genome to be indexed before execution. Similar to other aligners, we construct an index over the reference genome in a separate, offline phase. The index is a hash table of key-value pairs, where the key and value are both 32-bit unsigned integers. In order to populate the hash table, we extract k-mers from each position of the reference genome. As the reference sequence usually contains only 4 characters, namely A, T, C and G, we convert each character in the extracted k-mer into a two-bit equivalent representation. Any k-mers that contain ‘N’ characters are not added to the index. The k-mer length is a configurable parameter in Accel-Align, but we set it to 32 to enable a k-mer to fit in a single 64-bit integer. We hash the k-mer to generate the key by using a simple modulo-based hash function that maps the 64-bit integer into one of *M* buckets, where *M* is a large prime number that fits in a 32-bit integer. The 32-bit reference location offset from where the k-mer was extracted is the value associated with the key.

As the hash table is repeatedly used for looking up candidates during alignment, it is important to physically store these key–value pairs efficiently. We do this by using a chained hash table implementation based on two flat 32-bit integer arrays. With our construction, there are at most *M* different keys and *N − k* + 1 different values, where *N* is the length of reference genome. As multiple k-mers can hash to the same key, each key can correspond to multiple position values. We gather all position values for each key, sort them individually, and store all such sorted values together, in key order, in a single *position* array. We represent the keys implicitly by an offset in a separate 4GB *key* array, and in each key-array entry, we store the cumulative count of candidate positions for all keys smaller than that key. Thus, as the position array is ordered by key, all the candidate locations indexed between the offsets *K* and *K* + 1 in position table belong to the key *K*. The process is illustrated in Fig 1. Thus, the entire index, represented using the key and position arrays, is saved in a single file on disk, and loaded in memory whenever alignment starts.

**Fig 1.**
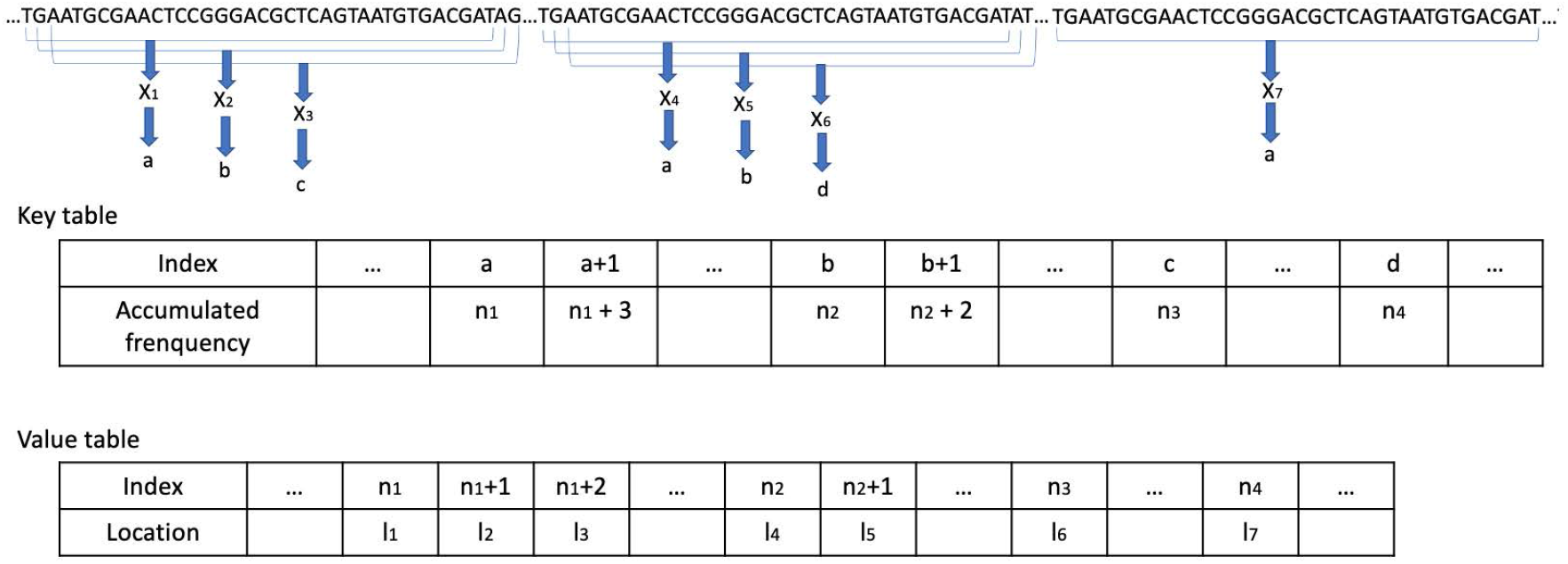
An index building example. *X*_*i*_ represents a 32-mer extracted from reference genome. *a, b, c, d* are the keys calculated from *X*_*i*_. *n*_*i*_ represents the number of keys smaller than the corresponding key. For example, *n*_1_is the number of keys smaller than *a*. Specifically, *X*_1_, *X*_4_ and *X*_7_ have the same key *a*. So there are *n*_1_ + 3 keys smaller than *a* + 1. Thus, *l*_1_, *l*_2_ and *l*_3_indexed with *n*_1_, *n*_1_+ 1, *n*_1_+ 2 in value table are the locations for *X*_1_, *X*_4_ and *X*_7_.

#### 2.1.2 Seeding

During seeding phase of alignment, we extract all non-overlapping k-mers of each read. For each k-mer, we compute the key, and use the key to extract the list of candidate positions. In situations where the seeds do not produce any candidates, we perform non-overlapping seeding after shifting the offset of the first seed by one position repeatedly until we find candidate positions. The positions are adjusted using the offset of the k-mer into the read to get normalized candidate positions. Then, we merge the candidate lists across k-mers to produce the final list of normalized positions that does not have any duplicates. One way of performing this merging is to gather all candidates in an array, sort it, and then find unique elements. However, such an approach would take *O*(*N*_*l*_ lg *N*_*l*_) time if *N*_*l*_ is the total number of candidates across all k-mers.

We avoid sorting by exploiting the physical organization of our hash table index. The position array of the hash table contains a key-ordered list of candidate positions, where each key’s candidates are stored in sorted order. Thus, during seeding, the candidates retrieved for each k-mer will also be in sorted order. We maintain a min-priority queue of size *N*_*k*_, where *N*_*k*_ = *readlength/k* is the number of k-mers in each read, and initialize it with the first candidate location of each k-mer. Then, we pop the minimum value from the queue and push the next candidate from the same k-mer as the popped one into the queue. We repeat the pop–and–push steps until the all candidates have been processed. This approach allows us to process all candidates in *O*(*N*_*l*_) time without sorting as long as the number of k-mers is small enough to keep the overhead of min-priority-queue negligible, which we found to be the case for short-read alignment using an empirical evaluation.

In the case of single-end reads, we don’t apply any filtering to the merged candidate lists. Thus, all candidates are passed to the embedding stage. For paired-end reads, however, we use a configurable pairwise-distance threshold for identifying candidates from one read that have a matching pair within the specified distance in the other read. All such candidate pairs are passed to the embedding stage.

### 2.2 Embedding

After candidate locations are identified by seeding, Accel-Align moves to embedding, the second stage of SEE. We extract strings of length equal to the read length from the reference genome at each candidate location. The goal of embedding is to transform these reference strings, and the query string which is the read, into different strings such that the edit distance between the original strings can be approximated using the Hamming distance between new strings. We have implemented two randomized embedding algorithms in Accel-Align.

#### 2.2.1 3*N* -embedding

The first algorithm was proposed by Chakraborty *et al*. (2016) who showed that given two strings *x, y* of length *N* taken from an alphabet ∑ such that *d*_*E*_ (*x, y*), the edit distance between *x* and *y*, is less than *K*, there exists an embedding function *f* : ∑^*N*^ → ∑ ^3*N*^, such that the distortion *D*(*x, y*) = *d*_*H*_ (*f* (*x*), *f* (*y*))*/d*_*E*_ (*x, y*) lies in [1, *O*(*K*)] with at least 0.99 probability, where *d*_*H*_ (*x, y*) is the Hamming distance between the embedded strings. In other words, Chakraborty *et al*. (2016) proposed a randomized algorithm that can embed strings of length *N* into strings of length 3*N* such that Hamming distance of embedded strings is at most square of the edit distance between original strings. Recent studies have demonstrated that this algorithm, which we henceforth refer to as *3N-embedding (3NE)*, works well in practice for performing edit similarity joins for even relatively large edit distances (Zhang and Zhang (2017)).

Accel-Align uses 3NE for embedding both reference strings and the read itself. Listing 1 shows the pseudo-code for the embedding algorithm. The input string is a DNA sequence of length *N* consisting of four possible characters (A,C,G,T). The output is an embedding string of length 3*N* consisting of the four characters and possibly multiple repeats of a pad character (P). In each iteration, the algorithm appends a character from the input string, or the pad if it runs out of the input string, to the output string. Then, it uses a random binary bit string to decide if the input index should be advanced. The net effect of this algorithm is that some input characters appear uniquely in the output string, while others are randomly repeated multiple times. Using the theory of simple random walks, Chakraborty *et al*. (2016) established that the randomization in this algorithm will result in strings that differ by a small edit distance converging quickly to produce embedded strings that have a small Hamming distance.

##### Algorithm 1 3*N* –embedding

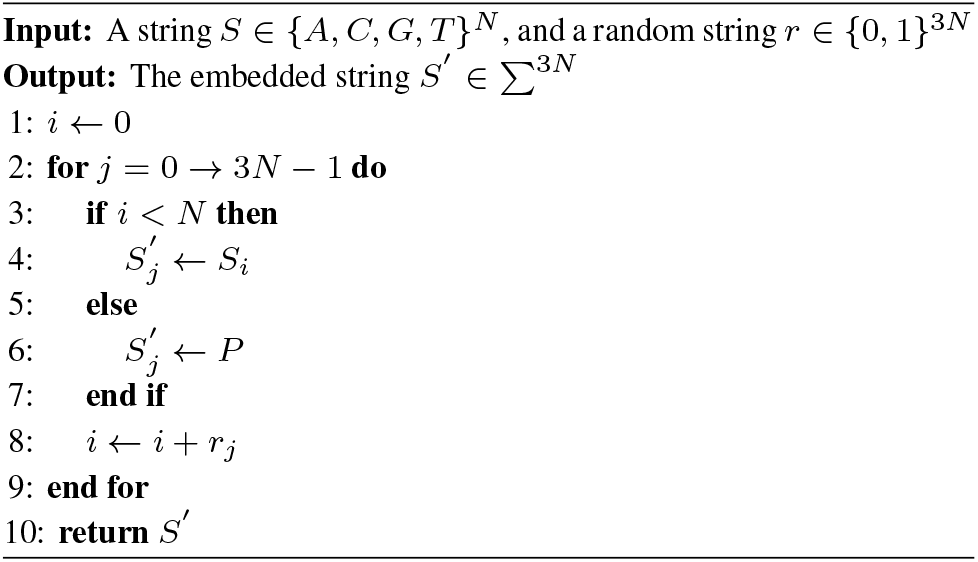

#### 2.2.2 2*N* -embedding

An initial implementation of 3NE in Accel-Align showed us early on that despite the simplicity of the algorithm, it was computationally intensive to embed billions of candidate locations across millions of reads. We describe several optimizations later in Section 2.4 that reduced the overhead of embedding, but one of the first optimizations we designed was a variant of the embedding proposed by Chakraborty *et al*. (2016), which we refer to as *2N-Embedding (2NE)*. Listing 2 shows the pseudocode for 2NE which is conceptually similar to 3NE with the exception that each character in the input string is copied to the output string at most two times. Thus, 2NE implements a mapping *f* : ∑^*N*^ *→* ∑^2*N*^ instead of ∑^*N*^ *→* ∑^3*N*^ . After implementing 2NE in Accel-Align, we found that Zhang *et al*. (2019) had also developed it in parallel, performed a theoretical analysis of its optimality, and used it for the edit similarity join application. As we show in our evaluation, we found 2NE to be functionally comparable to 3NE in terms of accuracy, and slightly better in terms of performance as it reduces the embedding time due to a reduction in the embedded string length from 3*N* to 2*N*.

##### Algorithm 2 2*N* –embedding

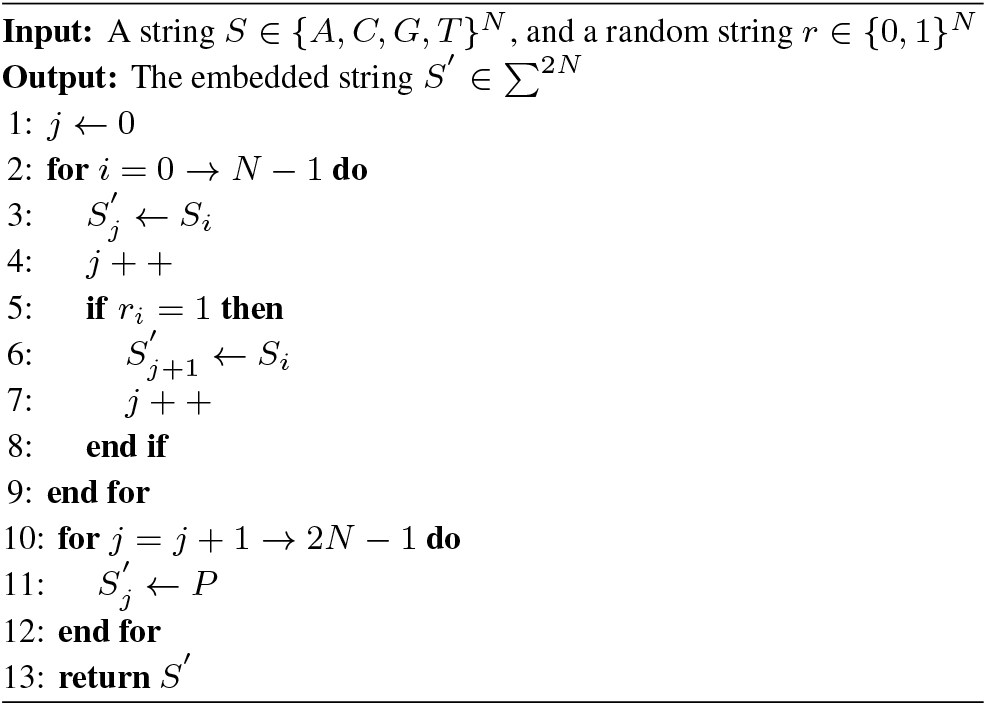

#### 2.2.3 Candidate Selection

We use one of the two embedding algorithms described above to embed all reference strings and the read. Then, we compute the Hamming distance between each embedded reference and the embedded read. We refer to this distance as the *embedding distance*. While doing so, we dynamically keep track of the top two candidate locations with least embedding distance and forward them to the third phase for further extension, scoring, and mapping quality computation.

### 2.3 Extension and MAPQ Computation

Before extension, Accel-Align will rectify the position of the best candidate. As mentioned in Section 2.1.2, the positions are adjusted using the offset of the k-mer into the read to get normalized candidate positions. However, this approach can shift the candidate position by a few nucleotides when there are indels. For example, consider a read with an insert in the first seed. If the second seed produces a candidate position *p*, then the expected normalized position is *p − k* + 1, not *p − k*. To fix this problem and identify the optimal position, we embed the first seed of the read and the k-mers in reference genome at multiple positions from *P − l* to *P* + *l*, where *P* is the best candidate position, *l* is the indel length threshold we set to 5% of the read length (this is conservative as the indel rate of short-read aligners is much lower). Finally, we take the starting position with the least embedding distance compared to the first seed of read as the final position.

Accel-Align can be configured to run in alignment-free mapping mode where only the identified candidate location is reported, or full-alignment mode where base-by-base extension is performed and the CIGAR string is reported. For the mapping mode, we simply pick the best candidate location, which is the one whose reference strings has the least embedding distance, and report its normalized location. For the full-alignment mode, we perform a global extension at the best candidate location using lib-ksw (Suzuki and Kasahara (2018)) with match(1), mismatch(4), gap-open(6), and gap-extension(1) penalties configured to match the settings used by BWA-MEM for computing the alignment score and CIGAR.

In addition to the CIGAR, popular aligners also report a mapping quality (MAPQ) that represents the degree of confidence in the alignment for each read. The basic idea behind MAPQ is that if multiple candidates align to the read with comparable alignment score, the aligner cannot be confident in its final choice. However, if alignment scores between the chosen candidate and others are very different, the aligner can be reasonably confident in its choice. Thus, MAPQ computation typically requires aligners to extend at least two candidate locations so that alignment scores can be compared. SEE, in contrast, makes it possible to avoid such extensions as we can exploit embedding distance once again to compute MAPQ. If two candidates are spaced apart with respect to edit distance from a read, then the embedded reference strings will also be spaced apart with respect to Hamming distance from the embedded read. Thus, Accel-Align uses Bowtie2’s MAPQ estimation procedure (Langmead and Salzberg (2012)) adjusted to use the embedded length, instead of the read length, and embedding distance of the top two candidates, instead of their alignment scores produced from Smith-Waterman extension, in order to produce a MAPQ value between 0 and 42.

### 2.4 Optimizations

A naive SEE implementation would perform seeding, embedding, and extension as described so far in sequence. However, during initial experimentation Accel-Align, we found that while embedding reduced the overhead of extension, the computational task of embedding and Hamming distance computation added non-negligible overhead. Thus, in addition to the 2NE algorithm described in Section 2.2.2, we implemented three other optimizations, namely, *pipelining, early-stop*, and *prioritizing*, that reduced the overhead of embedding without any change in functionality or accuracy.

#### Pipelining

Instead of embedding all candidates and then computing the Hamming distance, our first optimization is to pipeline these steps. We do this by modifying the embedding step so that the read is embedded first. Then each candidate location is embedded one by one, and the embedding algorithm simply updates the embedding distance in each iteration by comparing the output character of the candidate generated in that iteration with the corresponding character in the embedded read. This pipelining of embedding and distance computation provides three major benefits. First, as embedded strings are no longer generated in their entirety, it reduces memory consumption and associated overheads of allocating and freeing memory for storing embedded candidates. Second, it reduces the overhead caused by a needless second loop over the embedded candidates to calculate the Hamming distance. Third, as the output character generated by the algorithm is used immediately for distance computation, it improves processor cache utilization.

#### Early-stop

Using pipelining to produce the embedding distance for each candidate enables us to apply the second optimization based on the observation that only the top-two candidates with the least embedding distance are selected for further extension. Thus, if we have already encountered a candidate with a very low embedding distance, there is no point in continuing the embedding process for another candidate whose distance has already exceeded the previously observed minimum. Thus, we parameterize the embedding algorithm with a threshold such that the algorithm stops embedding a candidate as soon its embedding distance exceeds the threshold. Instead of storing the embedded distance of all candidates, we dynamically track the lowest and second-lowest distances, and simply use the latter as the threshold parameter.

#### Prioritizing

Our third optimization is a policy that drives pipelining and early-stop mechanisms. It is based on the intuition that if candidates with low embedding distance are prioritized before others, the overall cost of embedding will be low. This is due to the fact that the threshold will be set to a relatively low value during the early stages of embedding. As a result, early-stop will be applied to most candidates. However, the embedding distance of candidates is not known to us in advance. Therefore, we use candidate counting, a technique used by SFE aligners for count filtering (Liao *et al*. (2013b)), to prioritize candidates based on the assumption that candidates with higher counts or votes are more likely to have lower embedding distance, and more likely to be picked as the best candidate. Thus, we modify the seeding phase to associate with each candidate location a count of the number of k-mers that produced that location during the hash lookup. During embedding, we first embed the candidate with the highest count followed by all other candidates. It is important to note here that we still embed all candidates, albeit in a different order. Thus, unlike SFE aligners, we do not filter out candidates based on k-mer counting.

## 3 Results

In this section, we present an evaluation of Accel-Align using both simulated and real data to compare its performance and accuracy with respect to four state-of-the-art short-read aligners, namely, BWA-MEM (v0.7.17) (Li (2013)), Bowtie2 (v2.3.5) (Langmead and Salzberg (2012)), Minimap2 (v2.17) (Li (2018)), Subread (Liao *et al*. (2013a)). These aligners occupy different points on the design spectrum with respect to the choice of index (hash table versus FM index) and candidate filtering strategy (elimination versus selection) among other aspects. We also tried to evaluate SNAP (Zaharia *et al*. (2011)), but we do not report the results here as SNAP failed to index the reference on our server as it requires more memory than our server capacity. Accel-Align is implemented in C++ and is configured to use 2NE algorithm by default as it was found to be faster than 3NE with comparable accuracy as we demonstrate later. Accel-Align uses Intel Thread Building Blocks for parallelizing both index generation and alignment.

All experiments were run on a server equipped with a quad-core Intel(R) Core(TM) i5-7500 CPU clocked at 3.40GHz, 32GB RAM, and a 256GB SATA SSD. In an offline phase not reported here, we used each aligner to pre-index the reference genome. Then, in each alignment experiment, we run the aligner five times and gather execution statistics. As all aligners read the index from secondary storage, the first run is typically “cold” as data is not in memory. Table 1 shows the index size of each aligner, and time to load the reference and index with such a cold cache. As our index storage device is a SATA SSD with limited bandwidth (500 MB/s), aligners with larger indexes take longer to load the index. However, in practice, this is never an issue even with our commodity SSD, as a single run of any aligner results in the appropriate index being memory resident. Hence, in all remaining results, we ignore the first run and report performance of all aligners under “warm” execution when the index has already been cached in memory. As we found the “warm” performance of the last four runs to be stable with all aligners, we only report the average of last four execution times.

**Table 1.**
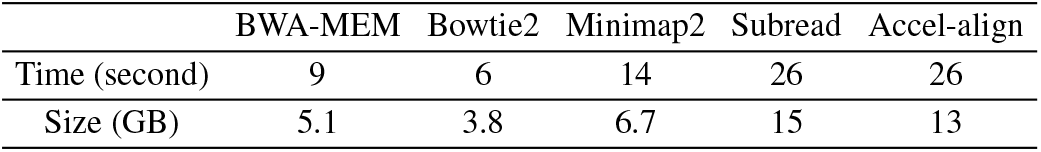
Time to load index

### 3.1 Benchmark with simulated short reads

For benchmarking Accel-Align, we used hg37 (hs37-1kg) as the reference genome. We used Mason2 (Holtgrewe (2010)) to generate a VCF file and then simulated reads from the VCF file together with an alignment file describing the exact coordinate of each read. We used Accel-Align, BWA-MEM, Bowtie2, Minimap2 (short-read mode), and Subread to align the reads and measured the end-to-end wall clock time for alignment. Using the Mason2 generated alignment file as our ground truth, we also evaluated the accuracy of each aligner with respect to two aspects. The first is the fraction of correctly mapped reads out of all input reads; we consider a read to be correctly mapped if the reported alignment overlaps with the Mason-provided one by at least 10% of read length. The second is the fraction of unmapped reads out of all input reads.

#### 3.1.1 Aligner comparison

Table 2 reports the performance (wall-clock execution time) and accuracy of the five aligners for a 10M, 100bp, single-end simulated read dataset generated by Mason2. In terms of performance, it can be seen that Accel-Align clearly outperforms the other aligners, as it is 9*×* faster than Bowtie2, 6*×* faster than BWA-MEM, 2.7*×* faster than Minimap2, and 5*×* faster than Subread.

**Table 2.** Evaluation on simulated single-end data (time format MM:SS)

|  | BWA-MEM | Bowtie2 | Minimap2 | Subread | Accel-align |
| --- | --- | --- | --- | --- | --- |
| Exec. time | 06:47 | 09:35 | 02:50 | 05:09 | 01:01 |
| Correctly mapped | 97.36% | 97.23% | 96.51% | 90.1% | 96.75% |
| Not mapped | 0.0028% | 0.044% | 0.73% | 9.64% | 0% |

Table 3 reports the performance and accuracy for a 10M, 100bp, paired-end dataset generated by Mason2. Accel-Align outperforms some aligners by an even larger margin here, as it is 11*×* faster than Bowtie2, 8*×* faster than BWA-MEM, 3*×* faster than Minimap2, and 4*×* faster than Subread. In terms of accuracy, Accel-Align is comparable with BWA-MEM, Bowtie2, and Minimap2, as a majority of reads are correctly mapped in both the single-end and paired-end datasets. Comparing Subread with BWA-MEM and Bowtie2, we can clearly see the performance–accuracy trade off of the voting strategy used by Subread for candidate filtering. Accel-Align, in contrast, can provide an order of magnitude improvement in performance over state-of-the-art aligners without sacrificing accuracy due to the use of embedding for candidate filtering.

**Table 3.** Evaluation on simulated paired-end data (time format MM:SS)

|  | BWA-MEM | Bowtie2 | Minimap2 | Subread | Accel-align |
| --- | --- | --- | --- | --- | --- |
| Exec. time | 14:49 | 19:37 | 04:58 | 06:32 | 01:40 |
| Correctly mapped | 98.54% | 98.51% | 98.09% | 92.39% | 98.24% |
| Not mapped | 0.002% | 0.0034% | 0.035% | 5.63% | 0.049% |

Fig 2 shows the RoC curve for the five aligners by plotting the fraction of reads mapped (Y-axis) and their error rate (X-axis) using each MAPQ value as a threshold (high to low from left to right). Once again, Subread has lowest accuracy at all MAPQ. Accel-Align generally performs better than Minimap2 but under performs BWA-MEM and Bowtie2 as it generally assigns fewer reads to higher MAPQ. Recall that Accel-Align, by default, uses embedding distance instead of alignment score for MAPQ computation. In order to isolate and understand the impact of embedding on MAPQ better, Fig 2 also shows the RoC curve for a modified version of Accel-Align, which we refer to as Accel-Align-AS, that uses embedding to pick the top two candidate locations and then uses alignment score obtained from extension of both candidates instead of the embedding distance to calculate the MAPQ.

**Fig 2.**
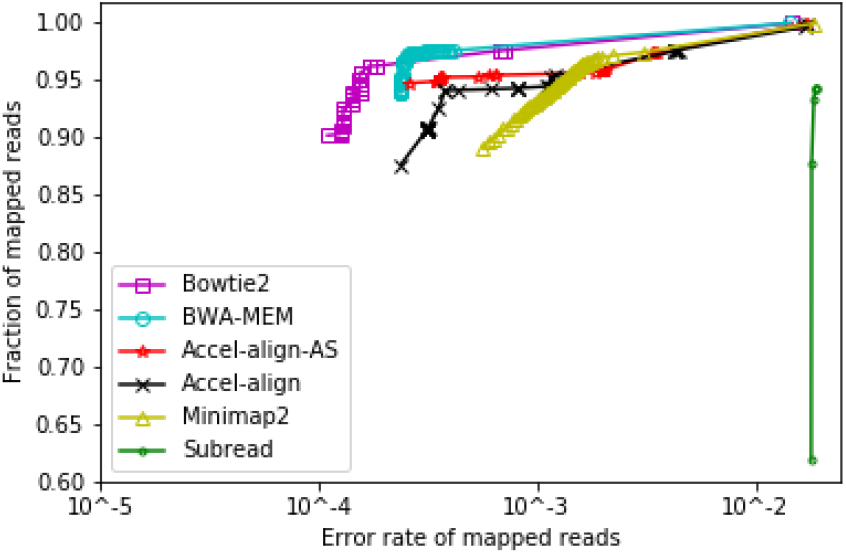
Error rate and fraction of reads over each MAPQ of 100bp pair-end reads.

Comparing Accel-Align with the Accel-Align-AS configuration, we see that Accel-Align-AS assigns a few more alignments (0.7%) to a higher MAPQ. As the only difference between these two configurations is the use of alignment score in Accel-Align-AS versus the embedding distance in the default version for computing MAPQ, we see that using embedding distance for MAPQ makes the aligner marginally more conservative in estimating MAPQ. Bowtie2’s MAPQ estimation procedure assigns MAPQ by comparing the best alignment score with respect to a threshold minimum score, and the second best score. We know that embedding distance computed for each candidate location is always larger than original edit distance. Thus, when embedding distance is used in MAPQ estimation, alignments that are further away from minimum with respect to alignment score end up moving closer to the minimum. This intuitively explains why MAPQ assignment is more conservative. However, it is important to note that the mapping location and difference in the overall error rate is the same for both Accel-Align and Accel-Align-AS. Thus, for the rest of this evaluation, we use the default version of Accel-Align.

#### 3.1.2 Varying read length

The computational cost of the embedding step is proportional to the read length, as each read and at least one candidate are converted from length *N* into a new string of length 2*N* . To test the sensitivity of performance with respect to read length, we used Mason2 to generate three more paired-end datasets with 10M reads, where each dataset was configured with a read length of either 125bp, 150bp, or 175bp. Table 4 shows the accuracy and Fig 3 shows the throughput (thousands of reads processed) per second per thread of the five aligners under various read lengths. The throughput of Bowtie2, Accel-Align and Subread is calculated by removing the reported reference and index preparation time from processing time. The throughput of BWA-MEM and Minimap2 is reported directly from their output log. Clearly, Accel-Align outperforms the other aligners at all read lengths as it provides 7–11*×* improvement over Bowtie2, 5–8*×* over BWA-MEM, 2–3*×* over Minimap2, 2-4*×* over Subread across a range of read lengths. Accel-Align correctly mapped around 0.2–0.5% less than BWA-MEM, Bowtie2 and Minimap2, while 1.6–5.8% more than Subread.

**Table 4.** Accuracy (% correctly mapped) on simulated paired-end data of different read length

|  | BWA-MEM | Bowtie2 | Minimap2 | Subread | Accel-align |
| --- | --- | --- | --- | --- | --- |
| 100bp | 98.54% | 98.51% | 98.09% | 92.39% | 98.24% |
| 125bp | 98.73% | 98.71% | 98.53% | 93.23% | 98.38% |
| 150bp | 98.82% | 98.80% | 98.72% | 93.65% | 98.41% |
| 175bp | 98.85% | 98.84% | 98.78% | 96.70% | 98.36% |

**Fig 3.**
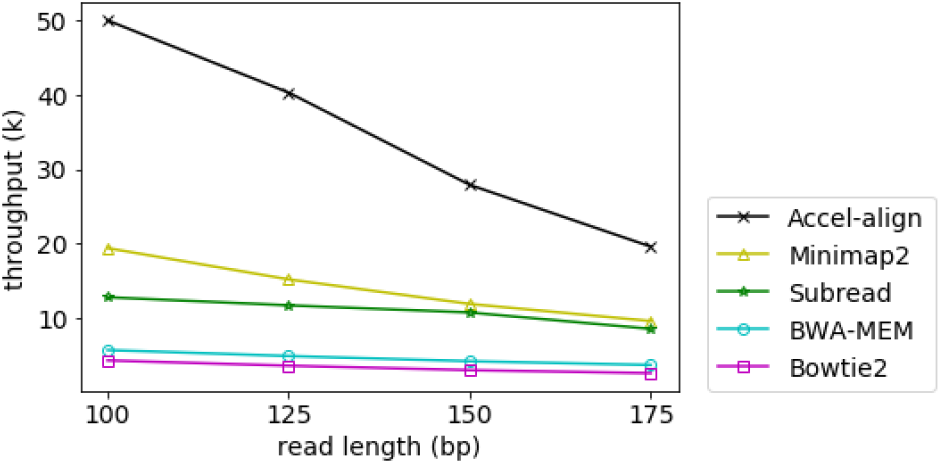
Throughput per second per thread for 100bp, 125bp, 150bp and 175bp pair-end reads.

#### 3.1.3 Alignment-free Mapping

Both Accel-Align and Minimap2 can be configured to run in alignment-free mapping mode where they report the position without the CIGAR string. The mapping mode completely eliminates the overhead of edit-distance computation. Although such mapping is useful in several applications that do not require base-by-base alignment, we use it in this context to isolate the benefit of embedding. To compare Accel-Align with Minimap2 in mapping mode, we used the two aligners to perform alignment-free mapping of the four paired-end Mason2 datasets. Fig 4 shows the execution time for various read lengths. Comparing Figs 3 and 4, we can make two observations. First, alignment-free mapping provides a further 1.3*×* improvement in throughput over base-to-base alignment. Accel-Align maps 10M, 100bp, paired-end reads in 75 seconds, thus, mapping more than 66,000 reads per second per thread without requiring any special hardware. These results demonstrate the benefit of using embedding in sequence mapping. Second, Accel-Align is 2.7– 3.5*×* faster than Minimap2 at all read lengths (Fig 4) with alignment-free mapping while offering compatible accuracy (Table 3). This demonstrates that our optimizations have eliminated any the computational overheads associated with embedding, making Accel-Align a competitive alternate to state-of-the-art sequence mappers.

**Fig 4.**
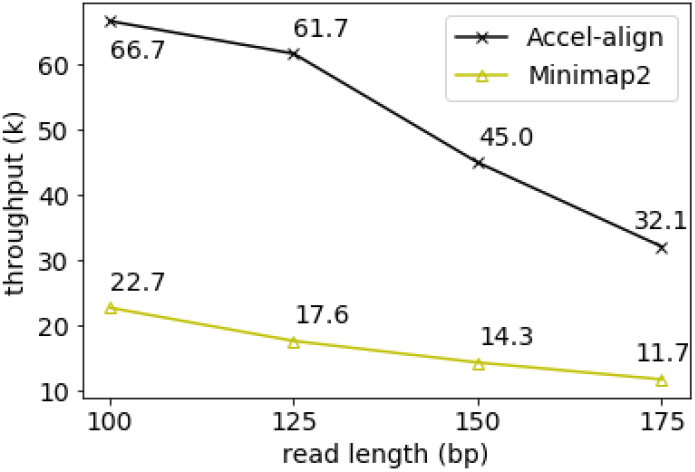
Alignment-free mapping throughput per second per thread for 100bp, 125bp, 150bp and 175bp read.

#### 3.1.4 Impact of embedding

To further isolate and understand the benefit of embedding in Accel-Align, we modified Accel-Align to run in three modes. (i) a *no-embed* mode where embedding is not used, and all candidate locations identified by seeding are directly forwarded for extension, (ii) the default mode using 2NE, and (iii) using 3NE instead of 2NE. Table 5 shows the performance and accuracy results for these three modes under the 10M, 150bp, Mason2 pair-end, simulated-read dataset.

**Table 5.** Comparison of 2N-embedding and 3N-embedding

|  | No-embedding | 2N-embedding | 3N-embedding |
| --- | --- | --- | --- |
| Exec. time (MM:SS) | 25:29 | 02:59 | 03:51 |
| Correctly mapped | 98.73% | 98.41% | 98.37% |
| Not mapped | 0.0073% | 0.0073% | 0.0073% |

Comparing 2NE and 3NE cases, we can see that 2NE provides a 23% improvement in performance with no discernible difference in accuracy. Comparing 2NE and the no embedding cases, we see that embedding provides a 8*×* reduction in execution time as it is able to identify the optimal candidate location without relying on edit distance computations at a marginal 0.3% lower accuracy.

Recall that Accel-Align uses lib-ksw for extension. However, embedding is orthogonal to the type of extension technique used. To understand the effectiveness of embedding in the presence of alternate, more efficient extension methods, we incorporated the recently published Wavefront Alignment Algorithm (lib-wfa) (Marco-Sola *et al*. (2020)) as an alternative to lib-ksw for computing the alignment score and CIGAR. We modified Accel-Align to run in four modes. In *NO-KSW* and *NO-WFA* modes, we do not use embedding and pass all candidates for extension with either lib-ksw or lib-wfa. In *2NE-KSW* and *2NE-WFA* modes, we use 2NE for embedding to select top two candidate locations, and pass them for extension with lib-ksw or lib-wfa. Table 6 shows the performance for these four modes under the 10M, 150bp, Mason2 pair-end, simulated-read dataset.

**Table 6.**
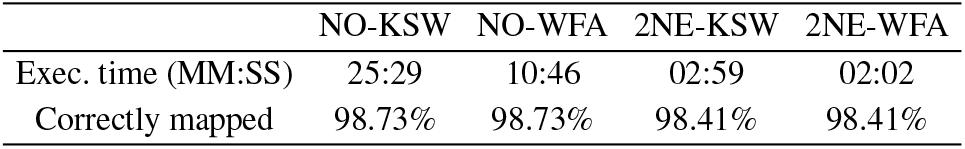
Comparison of optimized extension

|  | NO-KSW | NO-WFA | 2NE-KSW | 2NE-WFA |
| --- | --- | --- | --- | --- |
| Exec. time (MM:SS) | 25:29 | 10:46 | 02:59 | 02:02 |
| Correctly mapped | 98.73% | 98.73% | 98.41% | 98.41% |

There are several two important observations to be made from Table 6. First, we can see that replacing KSW with WFA provides a 2*×* reduction in execution time. 2NE-KSW provides another 4*×* reduction over NO-WFA due to the use of embedding. This shows that candidate filtering with embedding can provide a substantial improvement in performance even compared to optimized extension methods. Second, comparing 2NE-KSW and 2NE-WFA, we can see it is possible to achieve a further 1.4*×* improvement in execution time by combining embedding with optimized extension method like WFA. To maintain a fair comparison with other aligners which also use lib-ksw, particularly BWA-MEM and Minimap2, the rest of the results reported in this paper are based on lib-ksw.

### 3.2 Benchmark with real short reads

To evaluate the accuracy of Accel-Align on real data, we used the human whole-exome sequencing dataset NA12878 (accession No.: SRR098401). We built a pipeline similar to prior work Kumaran *et al*. (2019) to detect variants using GATK HaplotypeCaller (v4.1.0) (DePristo *et al*. (2011)). We used all aligners to align 85M paired-end reads in NA12878 to the hg37 reference genome. Then, we used the SureSelect Human All Exon v2 target captured kit bed file (ELID: S0293689) for capturing variant locations, and took high confidence variant calls (v2.19) from Genome in a Bottle (GiaB) consortium for validation.

We compare the aligners with respect to several metrics as shown in Table 7. The results for Accel-Align using the default parameters is shown in the column AA-32-mer. The execution time reports the wall-clock time taken by various aligners for aligning 85M paired-end reads (or 170M reads in total). Accel-Align provides a speedup of 9*×* over Bowtie2, 7.8*×* over BWA-MEM, 2.6*×* over Minimap2, and 4.8*×* over Subread, similar to the Mason2 dataset. The second metric is transition-to-transversion ratio (Ti/Tv), which is a key metric in detecting SNVs and should fall between 2.7–3.3 for this dataset, which is the case with all aligners. The last three metrics report precision, recall, and F-score values based on the GiaB truth set contains 23,686 SNVs and 1,258 indels contributing to a total of 24,944 variants for the NA12878 exome. The overall F-score of all four aligners are comparable except Subread. Accel-Align offers a slightly higher precision than the other aligners as most variants detected with Accel-Align are true variants. However, Accel-Align provides slightly lower recall, as it fails to detect some variants in the truth set.

**Table 7.** Evaluation on real data (time format HH:MM:SS)

|  | BWA-MEM | Bowtie2 | Minimap2 | Subread | AA-32-mer | AA-25-mer |
| --- | --- | --- | --- | --- | --- | --- |
| Exec. time | 1:32:02 | 1:48:09 | 00:31:24 | 00:56:24 | 00:11:46 | 00:19:20 |
| Ti/Tv | 2.71 | 2.73 | 2.73 | 2.85 | 2.78 | 2.76 |
| Precision | 0.961 | 0.967 | 0.967 | 0.980 | 0.972 | 0.968 |
| Recall | 0.952 | 0.948 | 0.952 | 0.901 | 0.942 | 0.946 |
| F-score | 0.956 | 0.957 | 0.959 | 0.939 | 0.957 | 0.958 |
| F-score(SNP) | 0.959 | 0.960 | 0.961 | 0.940 | 0.959 | 0.960 |
| F-score(InDels) | 0.920 | 0.915 | 0.927 | 0.914 | 0.912 | 0.915 |
| % Mapped | 99.86% | 99.16% | 99.55% | 91.43% | 96.92% | 98.67% |

To better understand the difference between detected variants, we show a Venn diagram of variants detected by various aligners in Fig 5. Clearly, 90.7% of variants are captured by all aligners. We find that there are 147 variants detected by the other four alignment tools, but not by Accel-Align. Upon further inspection, we found this to be due to two reasons. First, although embedding identifies the correct candidate location in most cases, there are reads for which it chooses the wrong candidate location. We found this to be particularly problematic for reads that map with a low edit distance to multiple locations in the reference. In such cases, we found that the hamming distance of embedded candidates is not spaced apart for embedding to identify a clear target. The second reason is the use of large k-mer length (32) in Accel-Align compared to the very short read length in NA12878 (76bp) which resulted in reads with two erroneous k-mers being unmapped. Table 7 shows the percent of mapped reads, and as can be seen, Accel-Align aligns 2.2-2.9% fewer reads comparing to BWA-MEM, Bowtie2 and Minimap2.

**Fig 5.**
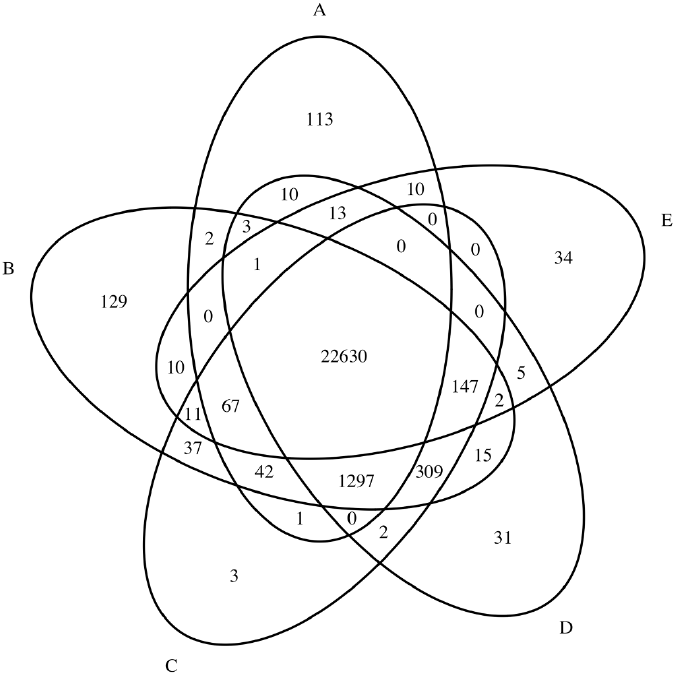
Venn diagram of variants detected by various aligners and Accel-Align (32-mer). A: Accel-align, B: BWA-MEM”, C: Minimap2, D: Bowtie2, E: Subread.

To understand the impact of k-mer size on overall accuracy, we modified Accel-Align to use 25-mers instead of 32-mers. Table 7 shows the results obtained using the 25-mer setting under column AA-25-mer. Comparing it with AA-32-mer, we have two important observations. First, the overall execution time increases by 39% compared to AA-32-mer case. Fig 6 shows a breakdown of execution time across the three stages when 25-mers or 32-mers are used. Clearly, this increase in time can be attributed almost entirely to seeding as 25-mers produce 4*×* more candidates than 32-mers. As SEE does not filter out any candidates, the overhead of candidate normalization, counting, and duplicate elimination performed during seeding increases. Despite this, embedding is able to identify the candidates, avoid needless extension, and still provide 2–6*×* reduction in execution time over other aligners. Second, the fraction of mapped reads increases by 1.7% by using 25-mers instead of 32-mers. Comparing Figs 5 and 7, we can see that this translates to an increase in the number of variants detected by Accel-Align. However, while the 25-mer case has better recall rate than the 32-mer case, it has a slightly lower precision as shown in Table 7. As a result, in terms of the overall F-score, the 25-mer case provides similar accuracy to the 32-mer case.

**Fig 6.**
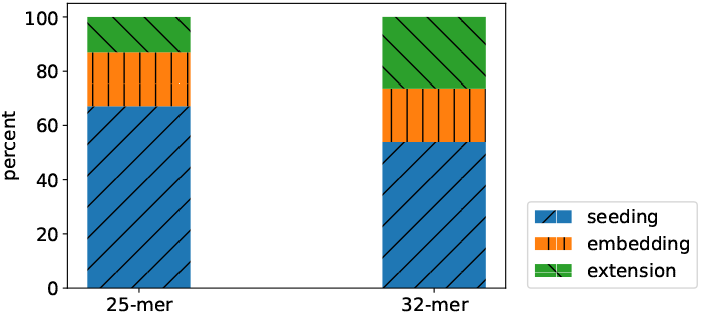
The percent of seeding, embedding and extension over the total processing time.

**Fig 7.**
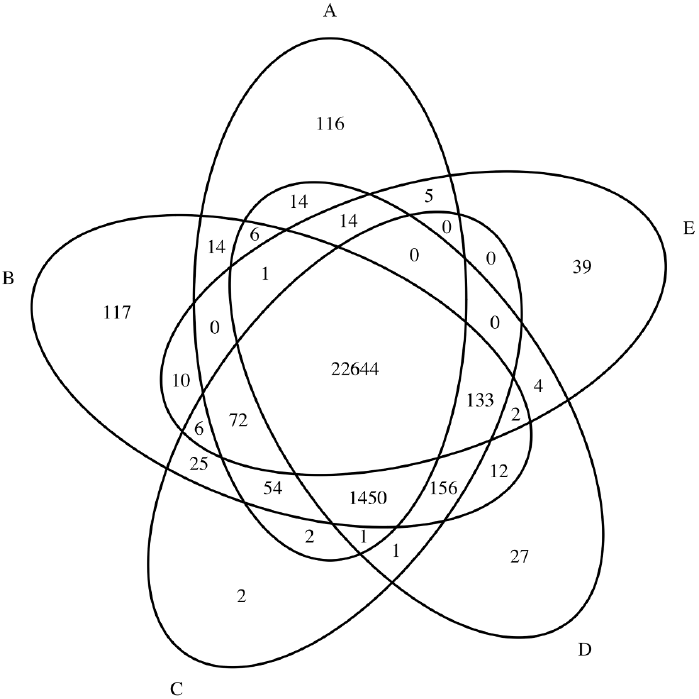
Venn diagram of variants detected by various aligners and Accel-Align (25-mer). A: Accel-align, B: BWA-MEM”, C: Minimap2, D: Bowtie2, E: Subread.

In order to isolate and understand the behavior of Accel-Align in the presence of SNP and indel variants, we show the F-score for SNP and InDels separately in Table 7 and show two Venn diagrams for the two types of variants separately in Fig 8. Clearly, Accel-Align is able to capture a majority of SNP and indel variants. In order to understand the impact of indel length on Accel-Align’s accuracy, we show the F-score of various aligners at each indel length in Fig 9. From these results we see that Accel-Align has the capability to detect even very long indels as evidenced by an F-score of 1 at the extremities in Fig 9. On investigation, we found that majority of indels missed by Accel-Align were short in length, and the main reason for lower accuracy was the inability of the non-overlapping seeding used by Accel-Align to produce the correct candidate location for embedding due to short 76nt read length of the NA12878 dataset. Our use of non-overlapping, sequential k-mers as seeds, and hash table as an index, was motivated by similar design choices made by other aligners (Zaharia *et al*. (2011); Marçais *et al*. (2018)) based on the assumption that with increasing read length and accuracy of short-read sequencers, it will be possible to draw enough, independent, long seeds with a high probability of at least one being error free. While researchers have demonstrated the benefit of other seed design strategies (like spaced-seeds (Ma *et al*. (2002)) and minimizers (Roberts *et al*. (2004))) and seed selection strategies (like cheap k-mer selection (Xin *et al*. (2013)), or dynamic programming (Xin *et al*. (2015a))), they are orthogonal to the problem of candidate filtering which is the focus of this paper. Similarly, in contrast to our fixed, but configurable, pair-wise filtering criteria, aligners also automatically learn the pair-filtering-distance dynamically, and use it to rescue paired reads where one of the pairs does not produce a mapping, or disambiguate otherwise equivalent mappings Canzar and Salzberg (2017). We leave open the task of combining embedding with better seed and pair selection strategies to future work.

**Fig 8.**
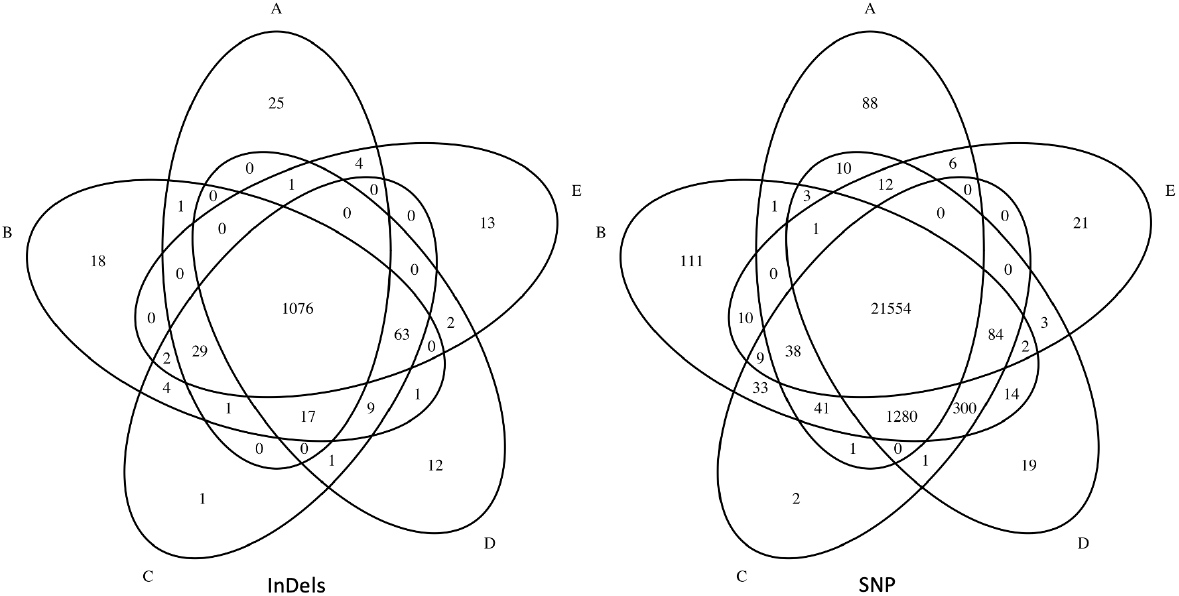
Venn diagram of SNP/InDels variants detected by various aligners and Accel-Align (32-mer). A: Accel-align, B: BWA-MEM”, C: Minimap2, D: Bowtie2, E: Subread.

**Fig 9.**
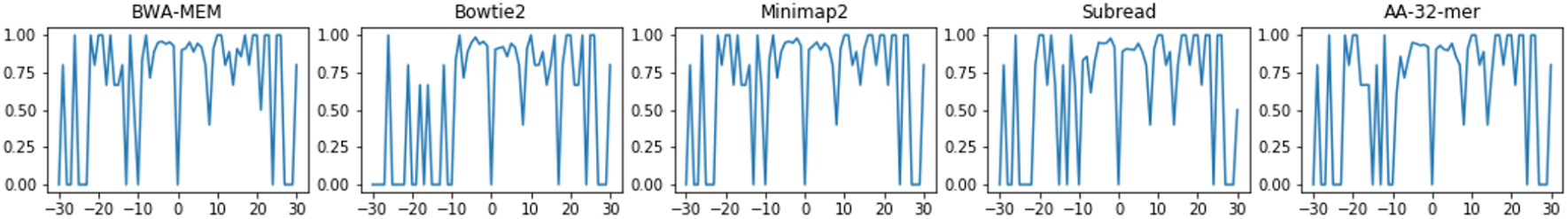
F-scores of InDels against the base pair length of the InDels.

## 4 Discussion

In this work, we presented Accel-Align–a fast sequence mapper and aligner that uses the SEE method for quickly identifying optimal candidates based on randomized embedding. Using an extensive evaluation, we showed that Accel-Align specifically, and SEE more generally, can provide an order of magnitude improvement over some state-of-the-art aligners on commodity CPUs. We are pursing several avenues of future work. First, we are working on extending Accel-Align to support additional functionalities like soft clipping and local alignment. Currently we do not support this as neither our current embedding approach, nor lib-ksw, support local extension.

In order to support soft clipping, Accel-Align cannot embed the entire read, as a position with a high embedding distance can still be the original mapping position chosen by local extension after soft clipping. Instead, we are investigating the use sub-sequences of a read between exactly-matched seeds as the embedding target.

Another natural extension of this work that we are pursuing is investigating the effectiveness of SEE for long-read alignment. Research on string-similarity join has demonstrated that these randomized embeddings excel when applied to long strings with even up to 20% mismatch (Zhang and Zhang (2017)). This suggests that SEE can also be used for aligning error-prone long reads generated by Nanopore or PacBio sequencers. Another line of work that we are exploring involves exploiting hardware-acceleration techniques for implementing SEE across CPUs and GPUs. Looking at Fig 6, we can see that seeding dominates overall execution time at low k-mer sizes and accounts for a substantial portion of time with large k-mers. In a recent microarchitectural study (Appuswamy *et al*. (2018)) of various sequence aligners, we demonstrated that the seeding stage is a prime candidate for acceleration by GPUs due to its data-parallel nature, unlike the extension stage which is difficult to parallelize due to data dependencies in dynamic programming. Thus, we are developing a parallel version of Accel-Align which can work across CPUs and GPUs and provide better accuracy, and a 10*×* further improvement in performance over the purely CPU-based Accel-Align.

## 5 Conclusions

As sequencing technologies continue to increase read length while improving throughput and accuracy, we believe that randomized embeddings open up new avenues for optimization that cannot be achieved by using edit distance. Accel-Align clearly demonstrates the potential of using embedding for short-read sequence alignment. We believe that the benefit of SEE methodology and low-distortion embedding are not limited to short-read alignment; any computational biology problem that is limited by the overhead of edit distance can benefit from embedding. As embedding transforms strings from edit to Hamming regime, computational tools like Locality Sensitive Hashing can also be applied on the resulting strings. Thus, the techniques presented in this paper have a much broader scope as they can be used for other applications like spliced RNA-seq and bi-sulfite alignment, multiple sequence alignment, and even sequence assembly.

